# Evaluation of Methodologies in Anti-nephrin Autoantibody Detection

**DOI:** 10.1101/2024.07.25.605154

**Authors:** Pan Liu, Shuping Liu, Vidhi Dalal, Jerome Lane, Elisa Gessaroli, Eleonora Forte, Lorenzo Gallon, Jing Jin

## Abstract

Recent studies discovered the prominent presence of anti-nephrin autoantibodies in minimal change disease, steroid-sensitive nephrotic syndrome and/or post-transplant recurrent focal segmental glomerulosclerosis (FSGS). However, widely different, and often unconventional autoantibody detection methods were used among these studies, making it challenging to assess the pathogenic role for the antibodies. Here we examined methods of conventional ELISA, magnetic on-beads ELISA, immunoprecipitation-immunoblotting (IP-IB), and cell- and tissue-based antibody assays with 127 plasma samples of kidney and non-kidney diseases. On the antigen side, we compared commercially available recombinant human nephrin extracelluar domain (ECD) produced from human or mouse cell lines, as well as lab-made full length, ECD, and series of ECD truncates for measuring autoantibody reactivity and specificity. Surprisingly, different assay methods and different antigen preparations led to observation of assay-specific false-positive and false-negative results. In general, a set of tests that combines magnetic beads-enhanced ELISA, followed by IP-IB, and epitope mapping showed the most robust results for anti-nephrin autoantibodies, detected in two primary FSGS patients among all cases tested. It is interesting to note that cell/tissue-based results, also supported by antigen truncation studies, clearly suggest steric hindrance of reactive epitopes, as in full length nephrin that forms compact self-associated complexes. In conclusion, anti-nephrin positivity is rare among the tested patients (2/127), including those with FSGS (2/42), and autoantibody results can be affected by the choice of detection methods.

## INTRODUCTION

Podocytopathy is associated with a broad spectrum of clinical conditions of podocyte effacement with disruption of the kidney filtration barrier. Besides genetic mutations, circulating factors injurious to the glomerulus have been proposed to cause some idiopathic kidney diseases^1, 2^, and plasmapheresis is partially effective to reverse proteinuria and achieve remission^3^. Ongoing investigation of the exact nature of the so-called “permeability factor”, particularly in the context of focal segmental glomerulosclerosis (FSGS) in adults and steroid-resistant nephrotic syndrome in children, remains an active area of research, and the recent discovery of anti-nephrin autoantibodies in FSGS, pediatric nephrotic syndrome, and minimal change disease (MCD) sheds light on the etiology of podocytopathies^4-9^. Importantly, it was recently reported a strong correlation of autoantibody levels with disease activity, and/or treatment response to immunosuppressants, further suggesting a direct causative role of anti-nephrin antibodies.

Despite anti-nephrin antibodies being measured in the patients’ plasma, kidney biopsies show a low degree of colocalization between the antibody and nephrin at the slit junctions of neighboring foot processes. Instead, IgGs are mostly seen in intracellular puncta with nephrin, implicating that the autoantibodies cause rapid internalization of nephrin molecules from cell surface. However, the lack of convincing colocalization patterns between the antibody and nephrin at the slit membrane constitutes a challenge for the description of autoantibodies^4, 5^. Moreover, it also seems that the autoantibody titers are generally low and the reliability of conventional immune assays such as ELISA for detecting nephrin autoantibodies remains controversial^8^. These caveats call for a comprehensive evaluation of antibody detection methods for better understanding of detection thresholds, reproducibility, and feasibility of implementation in clinical laboratory settings. We sought to compare methodologies including the selection of recombinant nephrin antigens, the setup of antibody assays, and in-solution versus on-cell/tissue antibody detection methods.

## METHODS

Detailed methods are provided in the Supplementary Appendix.

## RESULTS AND DISCUSSION

### Summary of FSGS patient cohort, and major discrepancies in autoantibody results using mouse versus human cell line-produced recombinant nephrin antigen

Nephrin (NPHS1) is a 1241 amino acid transmembrane protein (Figure 1a). Like in the previous studies, we initially used the extracellular domain of nephrin (NPHS1-ECD) as the template sequence of recombinant antigens. These include His-tagged ECD produced from a mouse myeloma cell line (as in Shirai et al.^5^), His-tagged ECD produced from human embryonic kidney cells (HEK293) (as in Watts and Hengel et al.^4, 8^), and FLAG-tagged ECD from HEK293 production (Figure 1a, Supplementary Figure S1).

**Figure 1.**
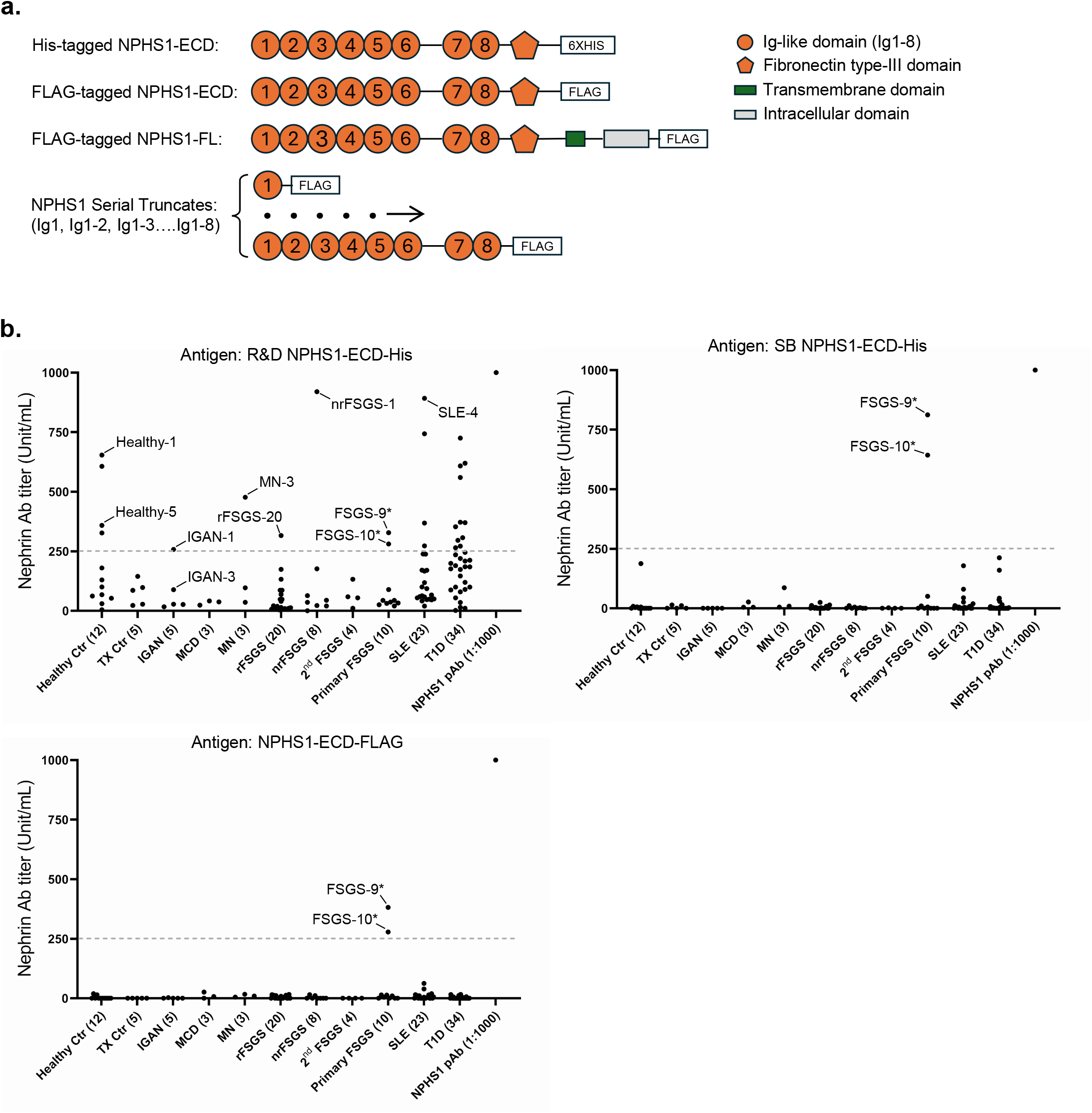
Discrepancies of ELISA result among different antigen preparations in anti-nephrin autoantibody detection. (**a**) Schematics of nephrin antigens used in the study. Various forms of NPHS1-ECD of the extracellular domain segment as antigen were used in ELISA in **b**. (**b**) Anti-NPHS1-ECD antibody titers measured by ELISA using different antigens, including commercial NPHS1-ECD from R&D Systems (produced in mouse cell lines) and Sino Biological (SB) (produced in HEK293 cells), and lab produced FLAG-tagged NPHS1-ECD (produced in HEK293 cells). Patient plasma samples were diluted 1: 300 in the R&D antigen group and 1:100 in other groups. The average signals from uncoated control wells were subtracted from the antigen-coated wells to eliminate non-specific background signals. The auto-antibody titers were calculated using a standard curve generated with serially diluted commercial NPHS1 polyclonal antibody, where the titer of 200 ng/mL polyclonal antibody was defined as 1000 U/mL. Note: in the SB antigen and NPHS1-ECD-FLAG ELISA groups, a subset of patients exhibited higher signals in uncoated wells compared to antigen-coated wells, resulting in negative values after subtraction. The titers of these samples were arbitrarily assigned a value of zero. Asterisks indicate samples recognized all the three antigens in ELISA.

With our focus on FSGS, the patient cohort was comprised of recurrent FSGS (rFSGS; n=20), nonrecurrent FSGS (nrFSGS; n=8), primary/idiopathic FSGS (n=10), and secondary FSGS (n=4), among a variety of additional kidney or non-kidney disease types^10^ (Supplementary Table S1). We also included a technical control using a commercial anti-nephrin antibody for standardizing antibody titer.

By coating the mouse versus human cell-produced nephrin ECD antigens (Supplementary Figure S1) to 96-well plates, we measured antibody activities in parallel by ELISA (Figure 1b). The results showed an alarming discrepancy between mouse versus human cell-produced NPHS1-ECD (His-tagged) (Figure 1b), with broad positive signals detected using the mouse cell-produced antigen, including from samples of healthy controls. The result was further confirmed by Western blotting using selected positive and negative samples against mouse cell-produced NPHS1-ECD (Supplementary Figure S2). This is in stark contrast to only two positive samples – depending on cutoff – reacting to HEK293-produced antigens, both from the primary FSGS group. These two cases, one being an adult-onset female and the other being a 17-months old boy, were denoted as FSGS-9 and FSGS-10, respectively. As murine cell lines can produce non-native glycosylation patterns, making recombinant proteins more immunogenic^11^,we then tested using PNGase F to remove *N*-glycans to rule out the reactivities to mouse-produced ECD attributable to species-specific glycoantigens. However, the PNGase F-treated antigen remained reactive to the false-positive plasma samples (Supplementary Figure S3a-b), and therefore we only used HEK293-produced recombinant nephrin antigens in all subsequent studies.

### Immunoprecipitation detection of anti-nephrin autoantibodies

One of the drawbacks of ELISA is that there is a wide variation of background signals among plasma samples that could render readings of negative values in antibody signals (as seen in Hengel et al.^8^). Therefore, we first tested several blocking buffers (a representative result is shown in Supplementary Figure S4). However, none of them effectively lowered backgrounds. Like in those prior studies^4, 6-8^, we alternatively conducted immunoprecipitation (IP) detection of antibody-nephrin interactions, developed in conjunction with subsequent Western blotting (WB) for visualizing the bound antigen. Consistent with the ELISA results, this IP-WB workflow confirmed the prominent presence of anti-nephrin antibodies in FSGS-9 and FSGS-10 samples (Figure 2a).

**Figure 2.**
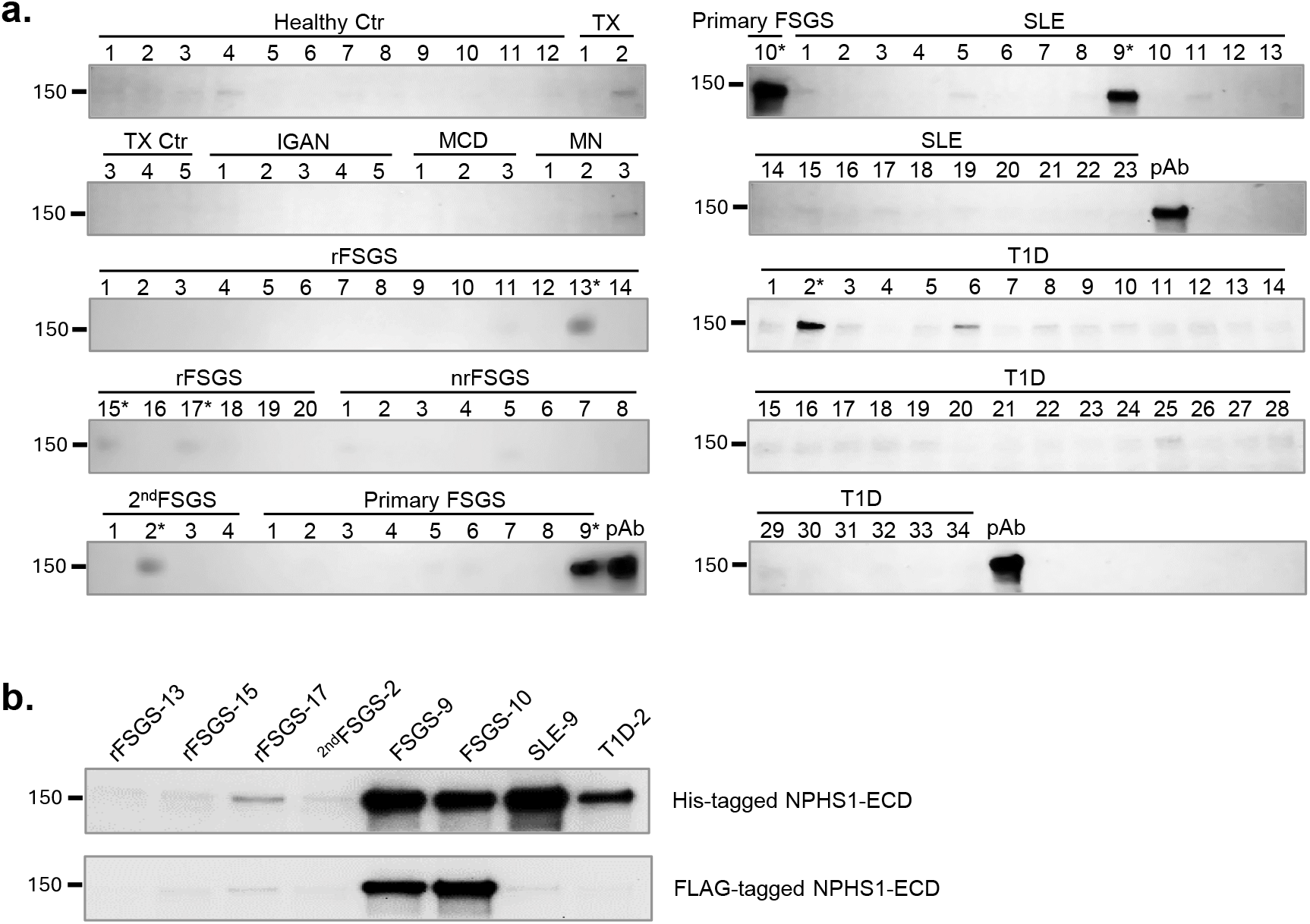
Detection of anti-nephrin autoantibodies by immunoprecipitation and Western blot using NPHS1-ECD antigens. (**a**) Western blot detection of NPHS1-ECD-His immunoprecipitated by either antibody control (pAb: sheep polycloncal anti-nephrin antibody) or patient plasma samples. To minimize variation caused by Western blot procedures, samples from different groups were blotted on the same membrane. SDS-PAGE gels were cut at the 75 kDa and above the 250 kDa protein marker positions. Gel slices containing samples from Healthy control, TX, IGAN, MCD, MN, primary FSGS-10, and SLE were transferred to one PVDF membrane. Gel slices with rFSGS, nrFSGS, ^2nd^FSGS, and primary FSGS 1-9 samples were transferred to a second PVDF membrane, while gel slices with T1D samples were transferred to a third PVDF membrane. To facilitate comparison across different membranes, a positive control of pAb immunoprecipitated NPHS1-ECD-His was included in each membrane. (**b**) Comparison of immunoprecipitation results for the selected positive plasma samples using NPHS1-ECD-His or NPHS1-ECD-FLAG as the antigen. A His vs. FLAG tag-directed discrepancy was evident in samples SLE-9 and T1D-2, whereas true anti-nephrin reactivities of FSGS-9 and FSGS-10 were confirmed.

Meanwhile, there were also samples clearly presented with notable but low IP-WB signals, including samples from control disease types, such as one SLE and one T1D samples. We then tested them together with other weakly positive samples alongside the positive FSGS samples against an alternative FLAG-tagged NPHS1-ECD antigen (without His-tag). The result showed these SLE and T1D samples only interacted with the His-tagged antigen (Figure 2b), suggesting tag-specific false positivity of some patients.

### An at scale 96-well workflow using in-well immunoprecipitation with magnetic beads followed by on-beads ELISA

With the correct use of NPHS1 antigen, the IP-WB assay produced more reliable antibody results than ELISA which had high background issues. However, IP-WB is laborious and often difficult to implement in clinical diagnostic settings that require throughput and automation. Therefore, we next focused on developing a new 96-well workflow that combines IP and ELISA. We adapted the general concept of enhanced on-beads ELISA, in which biotinylated NPHS1-ECD (also FLAG-tagged) was incubated with patient plasma. Antibody-antigen complexes were then captured by Protein G-conjugated magnetic beads. Using a magnetic separator, all IP and subsequent washing steps were performed in 96-wells, and bound NPHS1-ECD was detected with Streptavidin-HRP and TMB substrate (Figure 3a). This protocol resulted in antibody signals all above those from their corresponding control beads, indicating an improvement from conventional ELISA (compare Figure 3b to Figure 1b). Furthermore, the on-beads results were consistent with IP-WB, with the FSGS-9 and FSGS-10 samples having the highest signals (Figure 3c). It is important to note that magnetic beads-mediated antibody detection can be fully automated (such as using KingFisher Flex instrument, ThermoFisher), or alternatively adapted to chemiluminescence immunoassay (CLIA) that is more compatible with the practice in clinical laboratories^12^.

**Figure 3.**
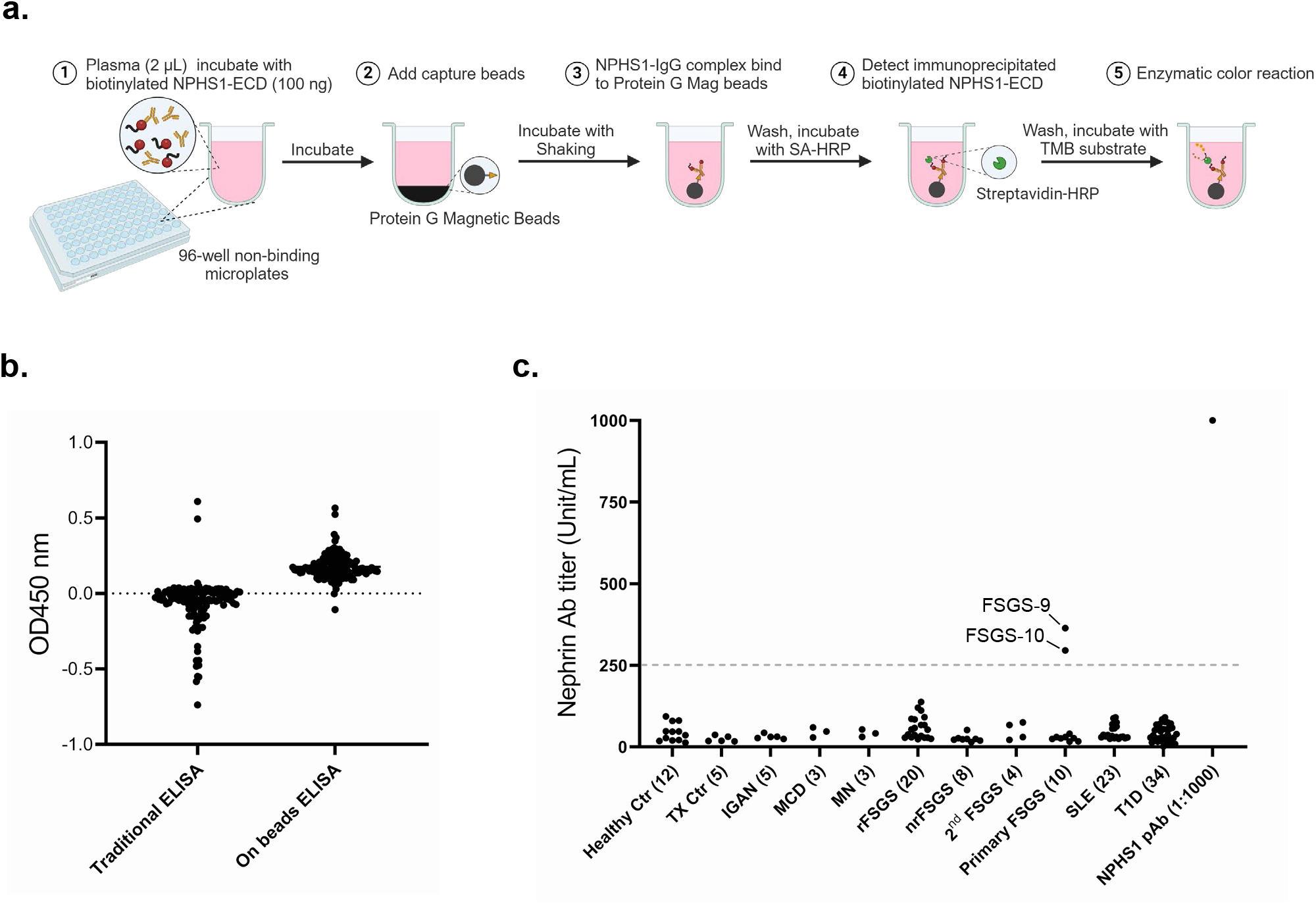
A high-throughput workflow for detection of NPHS1-ECD autoantibody by in-well IP and on-beads ELISA. (**a**) The general workflow resembles enhanced on-beads ELISA with the incorporation of in-well IP. (**b**) Comparison of autoantibody signal values (after subtracting background values from non-coated wells) between traditional ELISA and in-well IP followed by on-beads ELISA. This enhanced on-beads ELISA workflow demonstrated a significant advantage in eliminating non-specific background signals with the incorporation of a preceding in-well IP step. (**c**) Measurement of NPHS1-ECD antibody titers in all plasma samples using the Enhanced On-beads ELISA with biotinylated NPHS1-ECD-FLAG as the antigen.

### Epitope mapping via nephrin ECD truncation

To further investigate epitope-level reactivities of nephrin autoantibodies we constructed an additional series of nephrin truncates (Figure 4). Beside the full ECD sequence, we produced recombinant nephrin series of the first 1, 2, 3, 4, 5, 6, or 8 Ig-like domains (Figure 4a). These truncates were then tested with a negative plasma sample (Healthy-2) and the two positive samples (FSGS-9 and FSGS-10) in IP-WB. The Healthy-2 plasma did not recognize any truncates (Figure 4b). In contrast, the two positive samples showed distinct patterns of reactive signals to the truncates (Figure 4c, d), demonstrating nephrin epitopes could be patient-and/or disease-specific.

**Figure 4.**
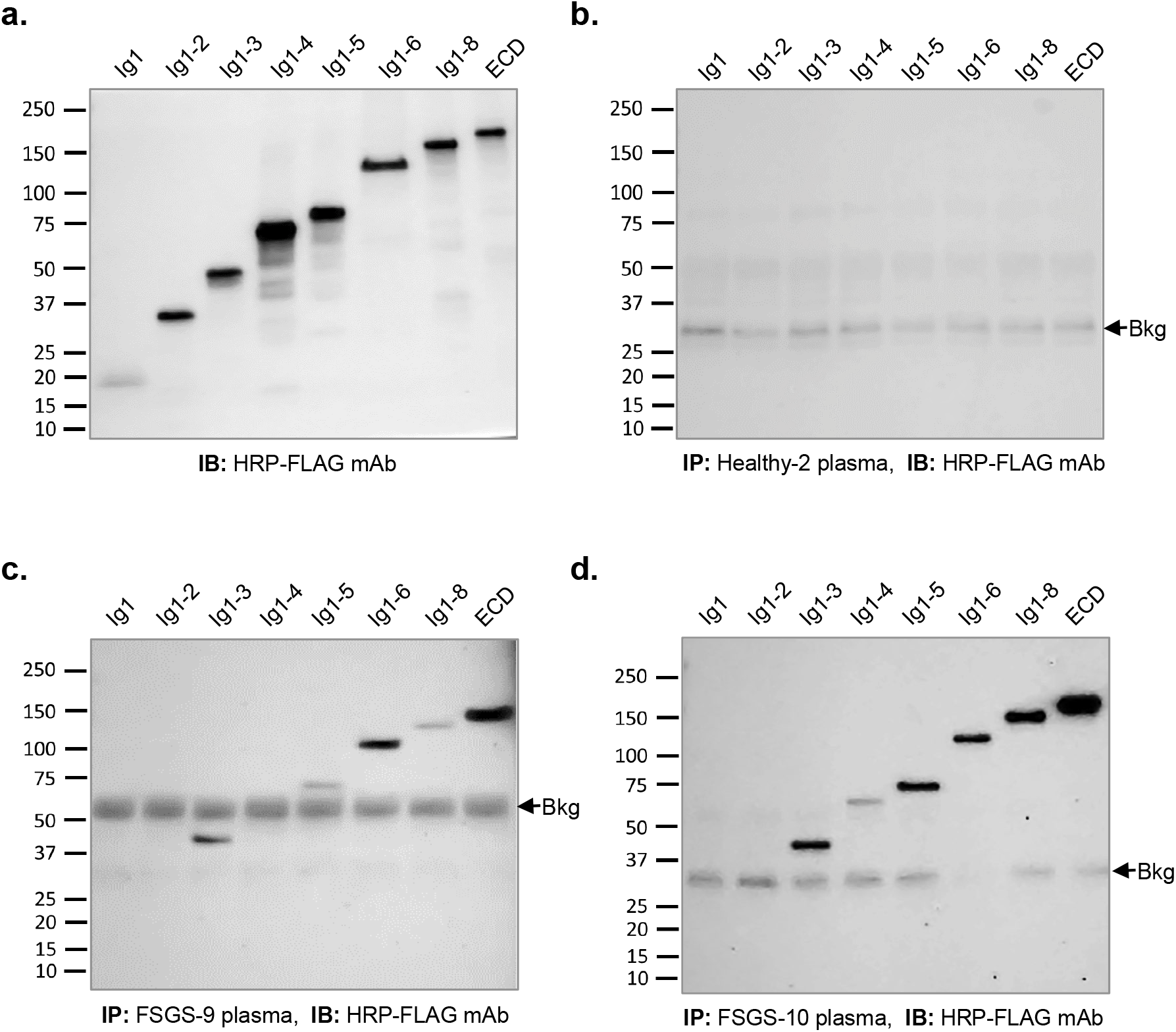
Epitope mapping using serial NPHS1 truncates in immunoprecipitation with healthy control and patient samples. (**a**) Western blot of recombinant NPHS1-ECD truncates using anti-FLAG tag antibody. (**b**) Western blot of the truncates immunoprecipitated with Healthy-2 negative control plasma. (**c**) Western blot of the truncates immunoprecipitated with FSGS-9 plasma. (**d**) Western blot of the truncates immunoprecipitated with FSGS-10 plasma, showing a different pattern of reactive signals across the truncates as compared to (c).

### Auto-antibodies only weakly react to oligomeric full-length nephrin

Nephrin is an integral component of the slit diaphragm through the formation high order ECD complexes at the junction of foot processes^13^. The self-association of ECD *in cis* is coordinated following interactions of the nephrin intracellular domain^14^. Indeed, when full-length nephrin was produced from HEK293 cells, it formed multimers (Figure 5a). This is in contrast to its ECD counterpart that was mostly monomeric. As expected, stable nephrin oligomers were maintained via intermolecular disulfide bridges^13^ (Figure 5a). Intriguingly, neither FSGS-9 nor FSGS-10 plasma reacted to this full-length nephrin oligomer (Figure 5b, Supplementary Figure S5, S6) in the IP-WB and ELISA assays, implicating possible steric hindrance of reactive epitopes in assembled nephrin structures.

**Figure 5.**
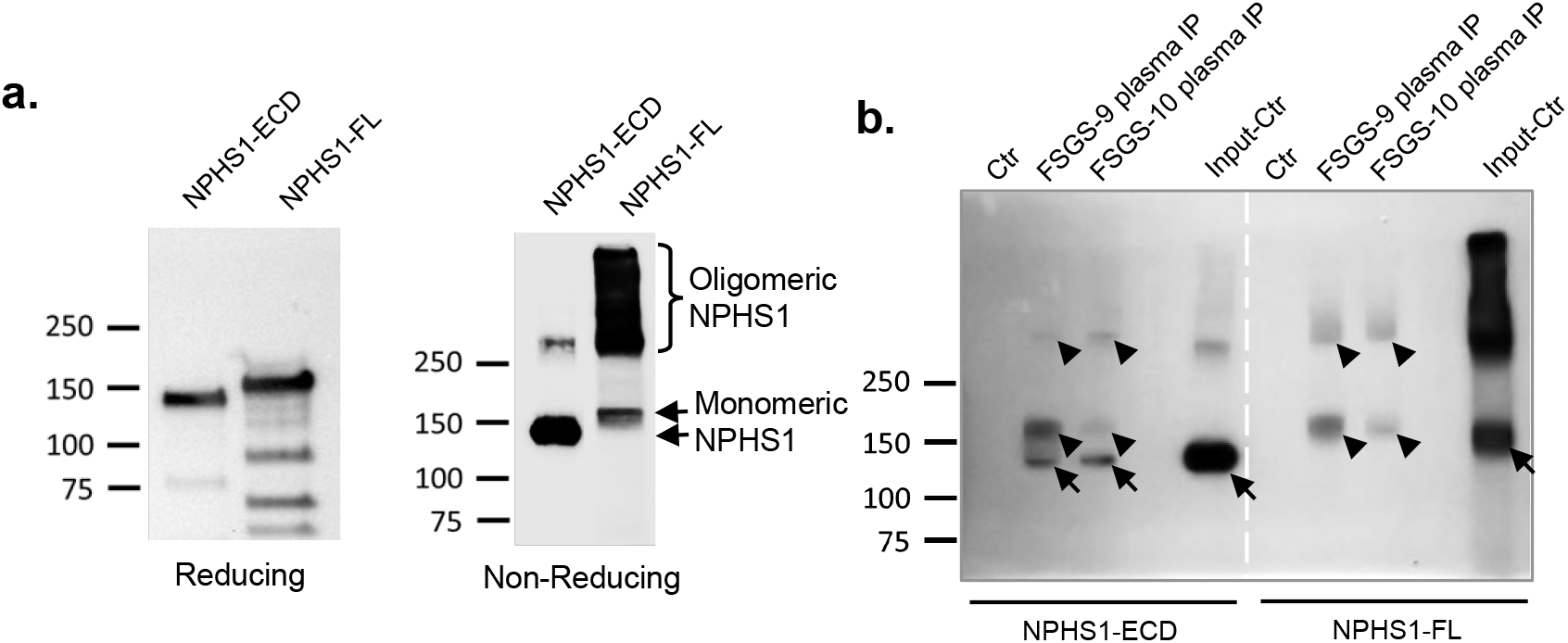
Full-length recombinant NPHS1-FL antigen reacts poorly to the anti-nephrin positive samples. (**a**) Western blot of recombinant NPHS1-ECD and full-length NPHS1-FL under reducing and non-reducing conditions. Both recombinant proteins were produced by clonal stable HEK293 cells. While NPHS1-ECD existed mostly as monomer, the full-length antigen was predominantly oligomers. (**b**) Western blots of NPHS1-ECD and full-length NPHS1-FL following immunoprecipitation by FSGS-9 and FSGS-10 plasma samples. The assay was performed under non-reducing conditions, and sheep anti-NPHS1 antibody was used to detect the recombinant proteins. Arrows indicate the monomeric NPHS1-ECD and NPHS1-FL bands. Arrowheads point to background bands of human immunoglobulins as contaminants, caused by a cross-reactivity with HRP-conjugated secondary antibody.

### Cell- and biopsy-based assays

To further investigate the lack of antibody reactivity to full-length nephrin, we transfected a plasmid expressing FLAG-tagged full-length nephrin in HEK293 cells. As expected, this protein was expressed at the cell surface, as detected using either anti-FLAG or anti-nephrin antibody by immunofluorescence (IF) (Figure 6a, b). However, directly probing the cell specimens with FSGS-9 and FSGS-10 plasma, in a fashion similar to that of anti-nuclear antibody assays^15^, failed to detect nephrin anchored at the cells surface (Figure 6a, b). Coincidentally, the lack of antibody signal in association with cell-anchored nephrin is somewhat similar to the low level of antibody presence in biopsy specimens together with nephrin at the slit diaphragm^5^.

**Figure 6.**
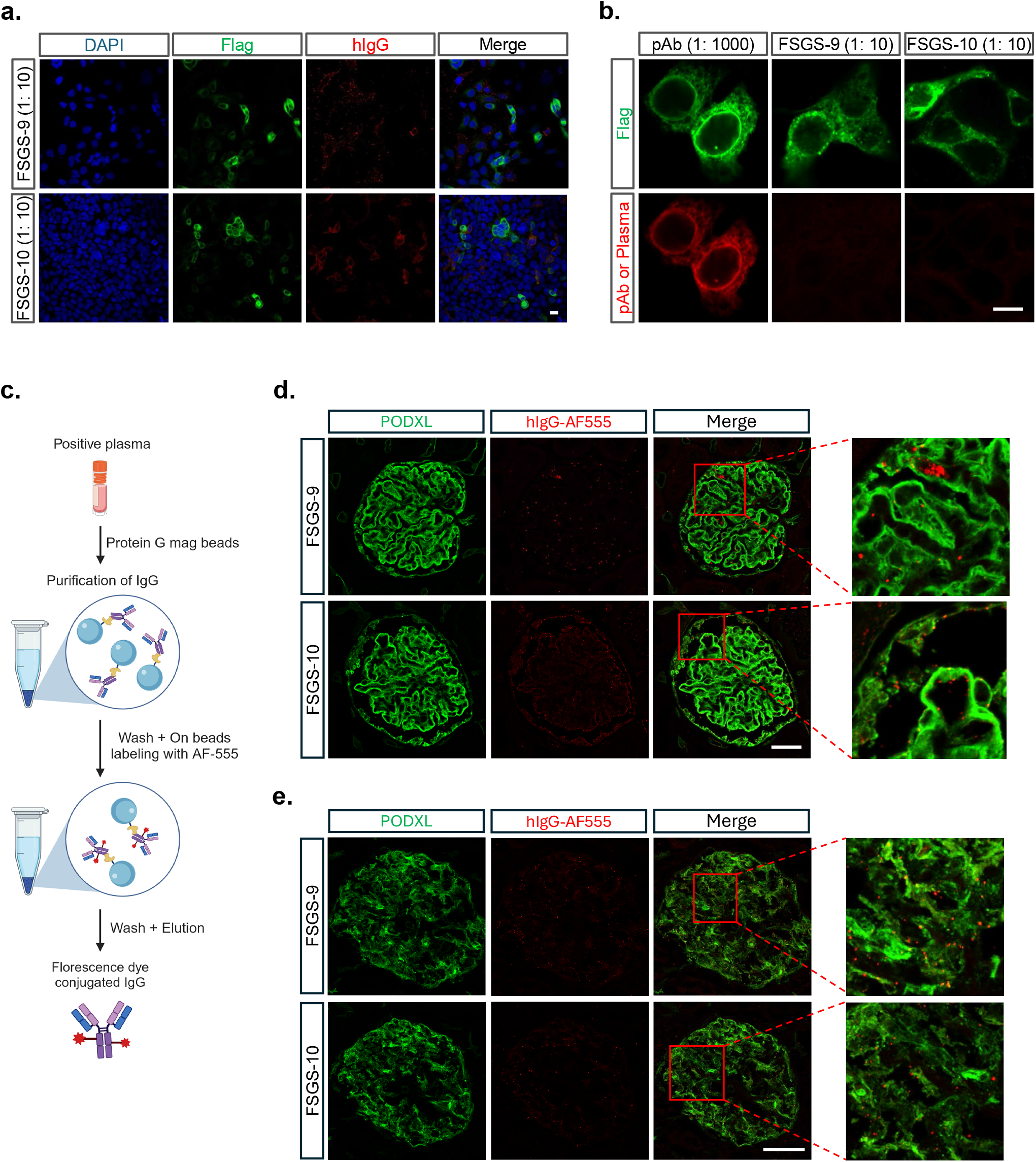
Anti-nephrin positive plasma showed no reactivity in both cell- and tissue-based assays. (**a**) Immunofluorescence staining of HEK293 cells transiently expressing FLAG-tagged full-length NPHS1-FL by FSGS-9 and FSGS-10 plasma (detecting human IgG in red, showing no overlaps with anti-FLAG counterstain in green). Scale bar: 10 μm. (**b**) Higher magnification of full-length NPHS1-FL at cell surface, showing no patient IgG staining of cell-expressed antigen. Scale bar: 10 μm. (**c**) Schematic representation of the process of on-beads AF-555 conjugation of IgG purified from positive plasma samples. (**d**-**e**) Immunofluorescence staining of paraffin-embedded human kidney sections (d) and frozen sections (e) using AF-555-labeled FSGS-9 and FSGS-10 IgG (in red). The podocyte slit diaphragm was counterstained for podocalyxin (PODXL in green) to mark the location of endogenous nephrin. No staining of the slit diaphragm was observed by patient IgG. Scale bar: 50 μm.

Similarly, we probed human kidney specimens using the two anti-nephrin positive plasma samples of FSGS-9 and FSGS-10 – with direct conjugation of patients’ IgG with fluorescence dye – to probe endogenous nephrin at the slit diaphragm. At comparable dilution as calculated from IP-WB signals between plasma and commercial anti-nephrin antibody, these patients’ IgGs failed to stain endogenous nephrin in both paraffin and frozen sections (Figure 6c-e, Supplementary Figure S7a-c). These results from cell- and tissue-based assays implicate the lack of antigen exposure to anti-nephrin autoantibodies in highly assembled slit structures, and that the hidden reactive epitopes are likely to be conformational.

## CONCLUSIONS

Immunoprecipitation-based methods provide robust detection of anti-nephrin autoantibodies, with which a high-throughput magnetic beads-directed assay is suitable for the first screening of serological activities. Recombinant nephrin of the extracellular domain that does not self-assemble into high-order oligomers reacts strongly with the autoantibody (Supplementary Figure S1). However, partly due to the low titer of anti-nephrin autoantibodies in plasma, the His-tagged NPHS1-ECD antigen is prone to tag-directed false positivity in some patients. Meanwhile, anti-nephrin autoantibodies do not generally react to endogenous nephrin that is densely assembled into high order structures at the slit diaphragm (A summary of all methods is in Supplementary Figure S8).

## DATA SHARING STATEMENT

All data generated through this study are available upon reasonable request to the Authors. Recombinant nephrin antigens can be made available to collaborators via a Material Transfer Agreement.

## ACKNOWLEDGEMENTS

We thank Dr. Anthony Chang of the University of Chicago for his insights on renal pathology. This work was partly supported through a grant from the National Institutes of Health (R01EB033377 to J.J.).

## AUTHOR CONTRIBUTIONS (requested in Style spreadsheet)

Conceptualization: P.L., J.J.

Methodology: P.L.

Investigation: P.L., S.L.

Formal Analysis: P.L., S.L., V.D., J.L., L.G., J.J.

Funding acquisition: L.G., J.J.

Manuscript preparation: All authors contributed.

