## Supplementary Appendix for "Evaluation of Methodologies in Anti-nephrin Autoantibody Detection"

#### Supplementary Material

- **Supplementary Methods**
- **Supplementary Figures S1-S8**
- **Supplementary Table 1**

#### SUPPLEMENTARY METHODS

##### Patient cohorts

Plasma samples were collected with patients' consent at Northwestern Memorial Hospital between 2014 and 2023 via IRBs STU00102396, STU00105534, STU00209738 and STU00208465.

##### Recombinant nephrin antigens

The various forms of human nephrin/NPHS1 proteins were either purchased from R&D Systems (6xHis-tagged extracellular domain of human nephrin – referred to as R&D NPHS1-ECD, aa23-1029 – produced using mouse myeloma cell line NS0, Cat# 93399-NN) and Sino Biological (referred to as SB NPHS1-ECD aa23-1055 produced using human-derived HEK293 cells, Cat# 17757-H08H), or produced in our own laboratory using cultured HEK293 and FreeStyle 293 cells, including full-length nephrin and the various truncates (see below).

To produce the Flag-tagged extracellular domain of human nephrin (NPHS1-ECD) protein, a DNA sequence encoding amino acids 1-1059 of NPHS1 with a C-terminal Flag tag (DYKDDDK) was synthesized by Integrated DNA Technologies. This cDNA was then ligated into the pcDNA3 vector (Invitrogen) for mammalian expression. The plasmid sequence was confirmed to be error-free through Sanger sequencing (ACGT Inc). Human embryonic kidney (HEK293) cells (ATCC) were transfected with the pcDNA3-NPHS1-ECD-Flag plasmid using TurboFect Transfection Reagent (ThermoFisher). After 24 hours, the transfected cells were serially diluted into 100 mm dishes and maintained in a selection medium supplemented with 1.2 mg/mL G418 (Thermo Fisher Scientific). The selection medium was replaced every three days until cell clones were visibly formed. These clonal cell masses were then individually seeded into new 48-well plates for amplification. After three days, the supernatants from each stable cell clone were tested for NPHS1-ECD-Flag expression using a standard ELISA method. An anti-NPHS1 polyclonal antibody (R&D System, Cat# MAB42692,) was used as the capture antibody, and a Horseradish Peroxidase (HRP)-conjugated Flag tag antibody (R&D System, Cat# HAM85291) was used as the detection antibody. To further confirm the stable clones with high NPHS1-ECD-Flag expression, the cell clones were cultured in FreeStyle 293 Expression Medium (ThermoFisher) for three days. The medium from each clone was then collected and precipitated using trichloroacetic acid (TCA). The precipitated protein pellets were washed with cold acetone and analyzed by western blot using the HRP-conjugated Flag tag antibody. The stable cell line with the highest expression level was amplified in DMEM complete medium (supplemented with 10% FBS, Penicillin-Streptomycin, and 1 mg/mL G418) until confluent and then maintained in FreeStyle 293 Expression Medium for five days. The conditioned FreeStyle 293 Expression Medium was then harvested for the purification of NPHS1-ECD-Flag recombinant protein. The protein was purified from the medium by ion exchange chromatography using HiTrap Q HP column (Cytiva, Cat# 17115401) followed by gel filtration using Superdex 200 Increase column (Cytiva, Cat# 28990944) with AKTA protein purification system (Cytiva). The purified protein was concentrated using Amicon Ultra Centrifugal Filter (50kDa MWCO, MilliporeSigma). To produce the Flag-tagged NPHS1-ECD truncates, the DNA fragments encoding aa23-142 for NPHS1-Ig1, aa23-241 for NPHS1-Ig1-2, aa23-339 for NPHS1-Ig1-3, aa23-439 for NPHS1-Ig1-4, aa23-543 for NPHS1-Ig1-5, aa23-739 for NPHS1-Ig1-6, aa23-939 for NPHS1-Ig1-8, were cloned into the pcDNA3 vector. The stable cell lines were generated, and the recombinant proteins were produced using the same protocol as for NPHS1-ECD-Flag production.

The full-length NPHS1 protein tagged with FLAG (NPHS1-FL-Flag) was produced in HEK293 cells using either transient transfection or stable cell line generation. Transiently transfected cells were lysed in NP-40 lysis buffer (50 mM Tris-HCl, 150 mM NaCl, 1% NP-40, and 5 mM EDTA, pH 7.4) supplemented with Protease Inhibitor Cocktail (MilliporeSigma). The cell lysate was clarified by centrifugation at 14,000 x g for 20 minutes at 4 °C. The NPHS1-FL-Flag protein was then purified using anti-FLAG M2 Affinity Gel (MilliporeSigma) following the manufacturer's instructions. The elution of the NPHS1-FL-Flag protein was achieved using 3x FLAG peptide (MilliporeSigma). The purified protein was concentrated using Amicon Ultra Centrifugal Filter which also removed excessive 3x FLAG peptide. For the stable cell line, cells were lysed in NP-40 lysis buffer, and the lysate was directly used for immunoprecipitation and western blot assays.

**ELISA**

ELISA microplates (96-well, Greiner Bio-One, Cat#655085) were coated with 100 ng/well of NPHS1-ECD-His, NPHS1-ECD-Flag, or NPHS1-FL-Flag overnight at 4 °C. All antigens were diluted in PBS buffer. In parallel, control plates were coated with PBS to measure non-specific binding background signals of the plasma samples. Following coating, plates were washed three times with

TBST buffer and blocked with blocking buffer for 1 hour at room temperature. Blocking buffers tested included 5% milk, 5% BSA, 1% fish gelatin, and SuperBlock blocking buffer (ThermoFisher, Cat#37515). The 5% milk in TBST showed optimal performance and was used in the following ELISA assays. After blocking, plates were incubated with plasma samples diluted in blocking buffer (1:300 for R&D NPHS1-ECD-His antigen, 1:100 for other NPHS1 antigens), and serial two-fold dilutions of sheep anti-hNPHS1 polyclonal antibodies were added to a row of wells on each plate to generate standard curves. After overnight incubation at 4 °C, plates were washed five times with TBST buffer and incubated for 2 hours with HRP-conjugated goat anti-human IgG Fc secondary antibody (1:4000, SouthernBiotech, Cat#2014-05) for wells containing human plasma samples, or HRP-conjugated donkey anti-sheep IgG secondary antibody (1:4000, R&D Systems, Cat#HAF016) for wells containing sheep anti-hNPHS1 polyclonal antibodies. After incubation, plates were washed five times with TBST and incubated with TMB substrate (BD Bioscience, Cat#555214). The reaction was stopped by adding 2N HCl when appropriate substrate color intensities were achieved. OD450 values were measured using a BioTek Microplate Reader. For each sample, the value from the uncoated well was subtracted from the coated well to account for non-specific binding. All ELISA assays were performed in triplicate.

Standard curves were generated based on the OD450 values from sheep anti-hNPHS1 polyclonal antibody wells. The titers of anti-NPHS1 autoantibody in plasma samples were calculated using these standard curves, with the titer of 1:1000 diluted sheep anti-hNPHS1 polyclonal antibody defined as 1000 U.

**Deglycosylation of NPHS1-ECD**

R&D NPHS1-ECD and SB NPHS1-ECD were treated with PNGase F Glycan Cleavage Kit (Gibco, Cat#A39245) to remove N-linked oligosaccharides under non-denaturing condition. 5 µg of each protein was mixed with 0.5 µL of PNGase F, and incubated at room temperature for 2 hours followed by overnight at 4 °C.

**Immunoprecipitation**

3 μL of human plasma sample, or 20 ng of sheep anti-NPHS1 polyclonal antibody as positive control was incubated with 100 ng NPHS1-ECD antigen, or cell lysate containing ~100 ng NPHS1-FL-Flag (quantified by ELISA) in 100 μL TBST overnight at 4 °C. When using cell lysate, protease inhibitor cocktail was added to the TBST to prevent potential antigen degradation. After incubation, 15 μL of pre-washed protein G magnetic beads (Cytiva, Cat# 28951379) was added to each tube and mixed for 4 hours at room temperature using a tube rotator (Labnet, Model H5500) to capture IgG-NPHS1 complex. The magnetic beads were then washed 5 times with TBST using DynaMa-2 Magnet (Invitrogen, Cat#12321D). The immune-complex was eluted with 15 μL Laemmli buffer (supplemented with or without 40 mM TCEP-HCl, Pierce, Cat# 20490) and incubated at 95 °C for 3 min.

**Western blot**

Protein samples were resolved on 4-15% pre-cast TGX gels (Bio-Rad, Cat# 4561085 or 4561085) and transferred to PVDF membranes using the Trans-Blot Turbo Kit (Bio-Rad, Cat# 1704274). After transfer, membranes were blocked with 5% milk for 1 hour at room temperature and then incubated overnight at 4 °C with sheep anti-NPHS1 polyclonal antibody (1: 800 diluted) or HRP-conjugated Rabbit anti-Flag tag antibody (1: 1000 diluted, R&D, Cat# HAM85291) in blocking buffer. When using the sheep anti-NPHS1 polyclonal antibody, the membranes were washed 3 times with TBST and then incubated with HRP-conjugated donkey anti-sheep IgG secondary antibody (R&D, Cat# HAF016) for 2 hours at room temperature. After 3 times wash with TBST, membranes were incubated with ECL Western Blotting Substrate (Promega, Cat# W1015) for 2 minutes and imaged using the iBright FL1500 Imaging System (Invitrogen, Cat# A44241).

**Biotinylation of Flag-tagged NPHS1-ECD protein**

Purified Flag-tagged NPHS1-ECD protein was labeled with EZ-Link™ NHS-LC-LC-Biotin (Thermo, Cat# 21343) according to the manufacturer's instructions. Briefly, 1 mg of Flag-tagged NPHS1-ECD dissolved in 250 μL PBS was mixed with EZ-Link NHS-LC-LC-Biotin at a molecular ratio of 1:3. The labeling reaction was carried out on ice for 3 hours. Excess biotin was then removed using Zeba Spin Desalting Columns (Thermo, Cat# 89890). The concentration of the biotinylated Flag-tagged NPHS1-ECD protein was adjusted to 1 mg/mL with PBS. The protein was aliquoted into small portions and stored in a -80°C freezer.

**96-well IP and on-beads ELISA**

96-well IP followed by on-beads ELISA assays were carried out in U-bottom 96-well non-binding microplates (Greiner, Cat# 650901). 2 µL of human plasma sample was incubated with 100 ng biotinylated FLAG-tagged NPHS1-ECD or vehicle control in 50 µL TBST overnight at 4 °C. Serial two-fold dilutions of sheep anti-hNPHS1 polyclonal antibody were used to generate standard curves. After incubation, 10 µL of protein G magnetic beads (pre-blocked with 5% BSA for 1 hour) were added to the wells and the plates were shaken at 400 rpm for 4 hours at room temperature using a microplate shaker (Fisher, Cat# 88861023). The beads were then washed 5 times with TBST using 96-well MagJET Separation Rack (Thermo, # FERMR03) and incubated with Streptavidin-HRP (1:2000 diluted in TBST, BD, Cat# 550946) at 400 rpm for 1 hour. The plates were washed 5 times with TBST, and incubated with TMB substrate until appropriate substrate color intensities were achieved. The reaction was stopped by adding 2N HCl, and the solution was transferred to new 96-well plates. OD450 values were measured using the same method described in ELISA section. For each sample, the value from the vehicle well was subtracted from the biotinylated Flag-tagged NPHS1-ECD well to account for non-specific binding. Standard curves were generated as described in ELISA section and used to calculate the titers of anti-NPHS1 autoantibody in plasma samples.

**On-bead fluorescence dye conjugation of IgG purified from plasma samples**

200 µL of plasma samples were diluted to 1 mL with TBST and incubated with 100 µL of protein G Mag Sepharose Xtra beads (Cytiva, Cat# 28967070) overnight at 4 °C with rotation. After incubation, the magnetic beads were washed three times with PBS and then resuspended in 0.1M $NaHCO_3$, 0.25M NaCl, pH 8.0. Subsequently, 4 µL of 10 mg/mL Alexa Fluor 555 NHS Ester (AF555, Invitrogen, Cat# A37571) dissolved in DMSO was added to each vial of beads and rotated for 2 hours at room temperature. The beads were again washed three times with PBS. AF555-labeled antibodies were then eluted twice with elution buffer (0.1M Glycine-HCl, pH 2.7) and immediately neutralized with 10 µL of neutralization buffer (1M Tris-HCl, pH 9.0), followed by concentration and buffer exchange using Amicon Ultra Centrifugal Filters. BSA and sodium azide were added to final concentrations of 0.1% and 0.02%, respectively. The titers of the labeled antibodies against NPHS1-ECD were measured by ELISA as previously described.

#### Immunofluorescence staining

For cell-based staining experiments, HEK293 cells were seeded in glass-bottom plates (Cellvis, Cat# P12-1.5H-N) pre-coated with 0.1% fish gelatin and transiently transfected with the plasmid encoding NPHS1-FL-Flag using TurboFect Transfection Reagent. Two days post-transfection, the cells were fixed with 4% PFA for 10 minutes, washed three times with TBST, and then blocked with blocking buffer (TBST containing 5% donkey serum and 0.1% saponin) for 1 hour. Primary antibodies or patient plasmas diluted in blocking buffer were incubated with the cells overnight at 4°C. After washing three times with washing buffer (TBST containing 0.1% saponin), secondary antibodies diluted in blocking buffer were added and incubated for 2 hours at room temperature. The nuclei were stained with DAPI for 10 minutes. The cells were washed 3 times again with washing buffer, and images were captured using a Nikon A1 confocal microscope.

For kidney sample-based staining, paraffin-embedded normal human kidney cortex sections (TissueArray, Cat# HuFPT072) and frozen normal human kidney sections (BioChain, Cat# T1234142) were used. For paraffin sections, after deparaffinization and rehydration, antigen unmasking was performed by incubating the slides in sub-boiled retrieval buffer (10 mM Sodium Citrate, pH 6.0) for 30 minutes. The subsequent blocking, primary-secondary antibody incubation, and image acquisition protocols were the same as those used for cell-based staining, except that the slides were mounted with Fluoromount-G (SouthernBiotech, Cat# 010001) before imaging.

### SUPPLEMENTARY FIGURE S1

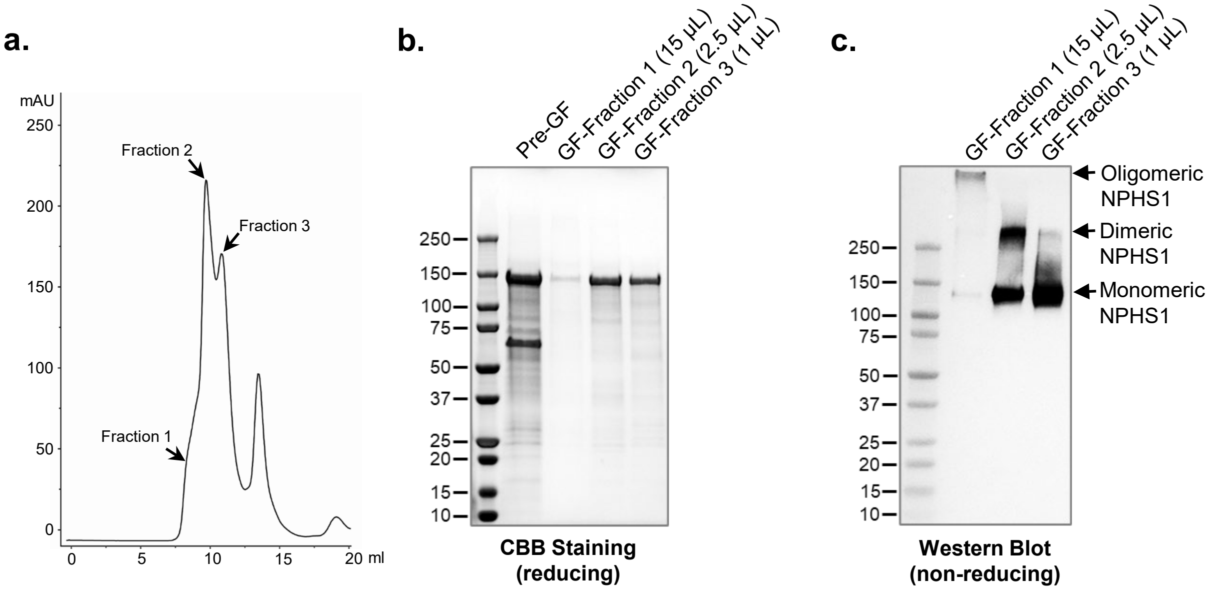


**Supplementary Figure S1. Production and gel filtration analysis of recombinant FLAG-tagged NPHS1-ECD.** Conditioned serum-free medium from stable HEK293 cell line expressing FLAG-tagged NPHS1-ECD was firstly purified through anion ion exchange chromatography, the fraction containing the recombinant protein was further purified by gel filtration. (**a**) Gel filtration (Superdex 200 increase) chromatography reveals the presence of multiple oligomeric forms of the recombinant FLAG-tagged NPHS1-ECD protein. (**b**) Coomassie blue staining of fractions collected from gel filtration chromatography confirmed all the 3 fractions are FLAG-tagged NPHS1-ECD. Samples loading volumes: fraction 1: fraction 2: fraction 3 = 15 : 2.5 : 1. (**c**) Western blot of the 3 fractions under non-reducing condition using anti-FLAG tag antibody. Loading volumes of each sample was the same as in (b). The results indicate that FLAG-tagged NPHS1-ECD primarily exists in a monomeric form, although small amounts of dimers and higher-order oligomers are also present.

### SUPPLEMENTARY FIGURE S2

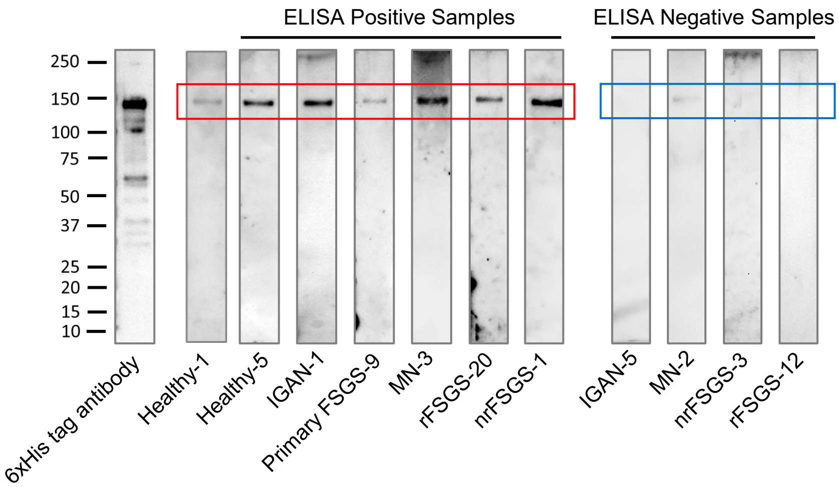


**Supplementary Figure S2. Western blot analysis confirmed the ELISA results for the R&D antigen.** Selected positive and negative plasma samples from the R&D 6xHis-tagged antigen ELISA assay were tested by western blot. The R&D antigen was resolved in 4-15% gels at 100 ng/well and then transferred to a PVDF membrane. The membranes was cut into strips and blotted with plasma diluted in blocking buffer (1:100), followed by detection with an HRP-conjugated anti-human IgG antibody. Membrane strip blotted with anti-His tag antibody was used as positive control. The red frame indicates that the R&D antigen was recognized by positive samples, while the blue frame indicates that negative samples did not recognize or only weakly recognized the R&D antigen.

### SUPPLEMENTARY FIGURE S3

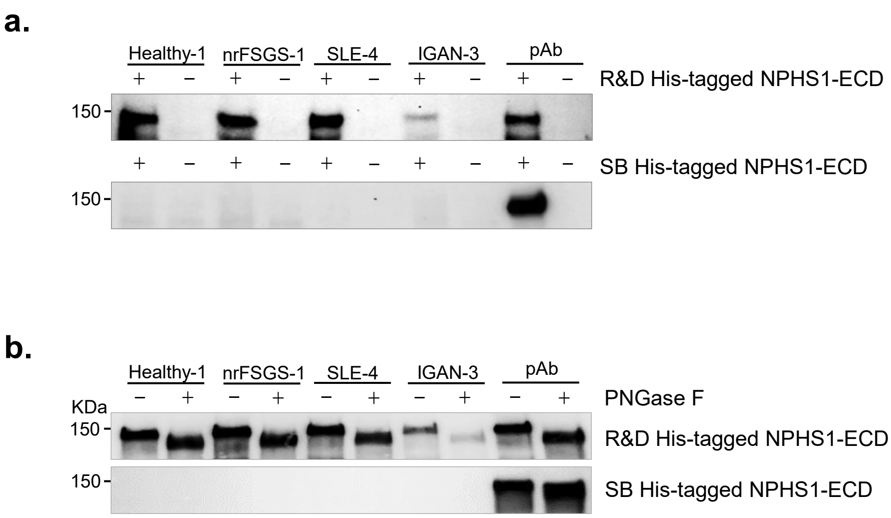


**Supplementary Figure S3. Immunoprecipitation analysis of selected plasma samples using R&D and SB antigens**. (**a**) Selected positive plasma samples recognize the R&D antigen but not the SB antigen in the immunoprecipitation assay. (**b**) Deglycosylation of antigens by PNGase F treatment did not change the antigen recognition pattern by plasma samples for both R&D and SB antigens.

### SUPPLEMENTARY FIGURE S4

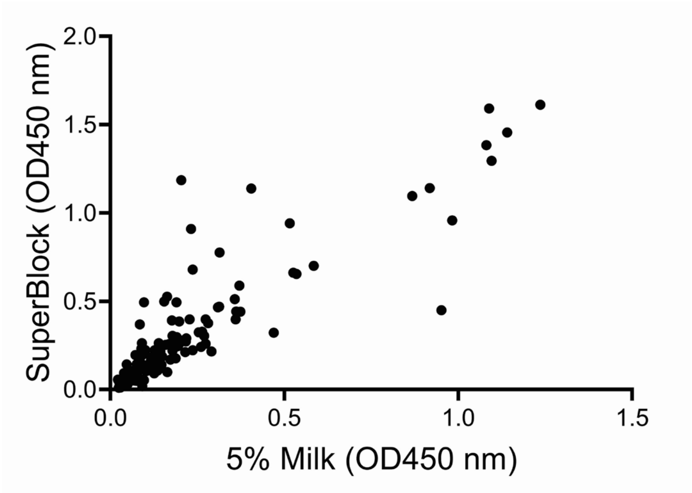


**Supplementary Figure S4.** Comparison of background signals of plasma samples in uncoated ELISA plate wells with SuperBlock and 5% milk blocking buffer.

### SUPPLEMENTARY FIGURE S5

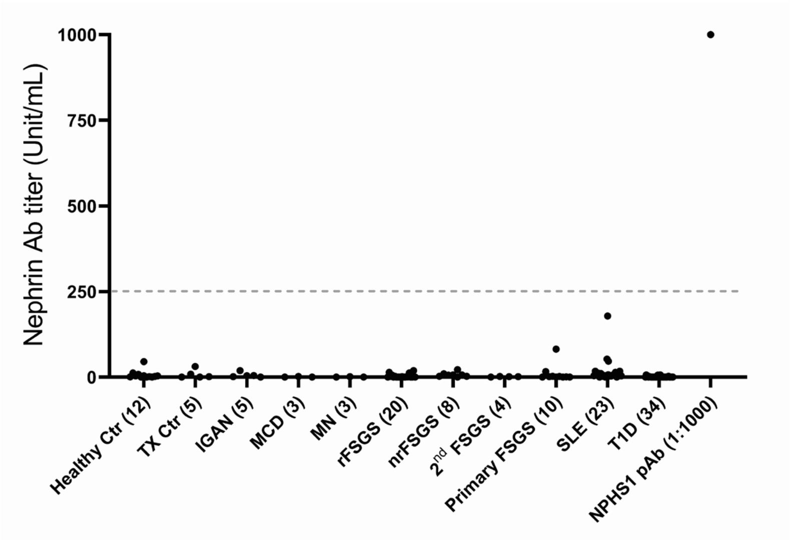


**Supplementary Figure S5.** Detection of NPHS1 autoantibody by ELISA using FLAG-tagged full-length NPHS1 protein. No positive sample was detected.

### SUPPLEMENTARY FIGURE S6

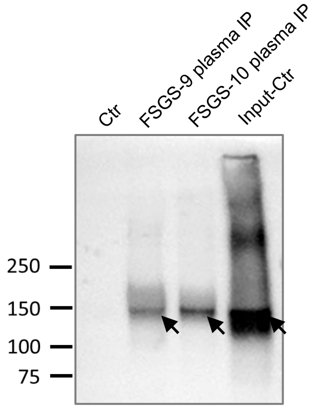


**Supplementary Figure S6. Immunoprecipitation of FLAG-tagged full-length NPHS1 produced by transient transfection.** Western blot was performed under non-reducing conditions. Compared to stable cell line expression (as in Figure 5), the transiently expressed NPHS1 shows a dominant monomeric form (arrows), indicating insufficient inter-molecular disulfide bond formation. The non-disulfide bond-mediated oligomers may be dissociated in the cell lysis buffer containing detergent, resulting in monomers that can be recognized by positive plasma FSGS-9 and FSGS-10.

### SUPPLEMENTARY FIGURE S7

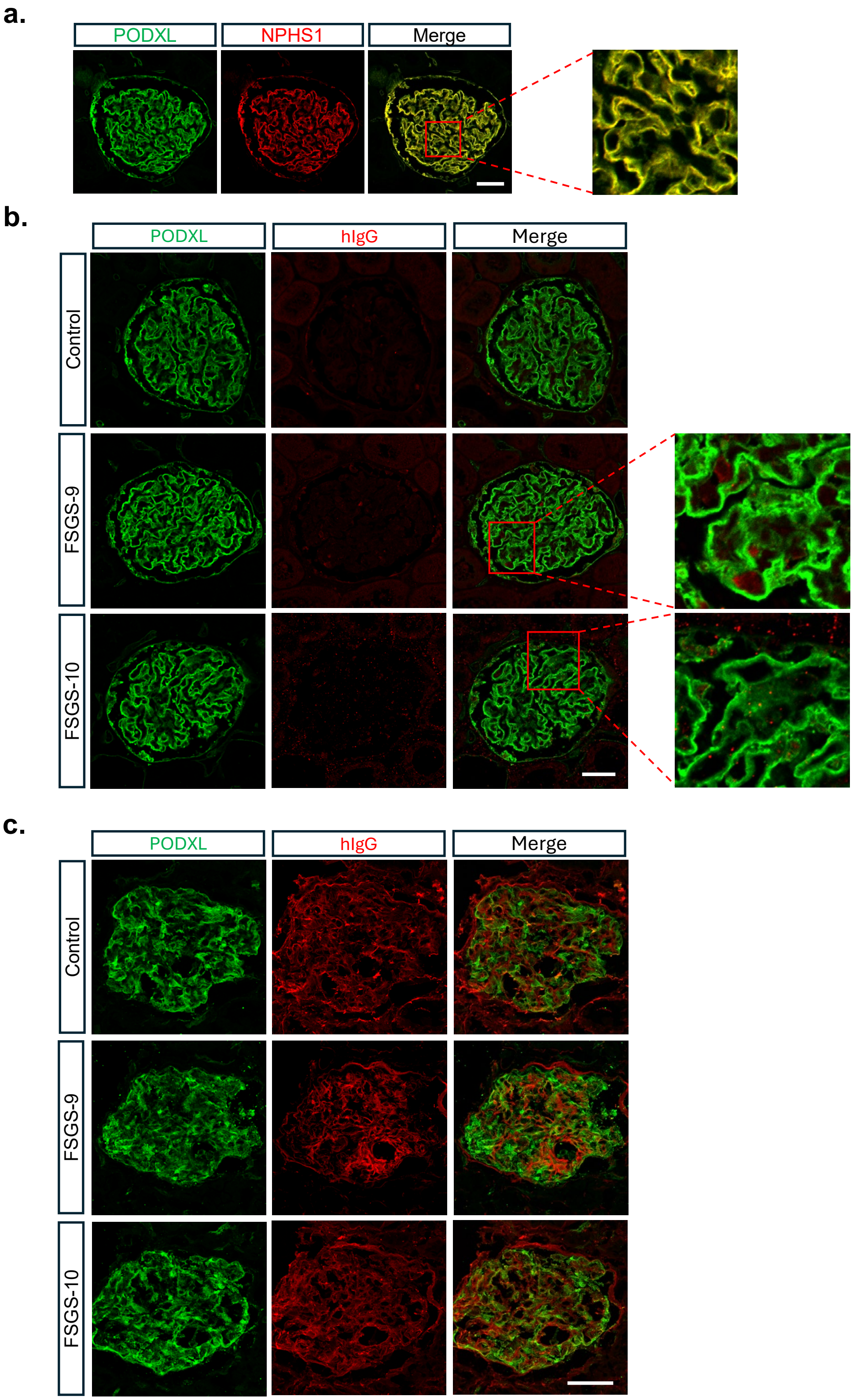


**Supplementary Figure S7. Immunofluorescence staining of normal human kidney sections with positive plasma samples.** (**a**) PODXL and NPHS1 exhibit consistent fluorescence localization in glomeruli; therefore, PODXL signal is used to represent NPHS1 distribution in glomeruli. (**b-c**) Immunofluorescence staining of paraffin-embedded normal human kidney sections (b) and frozen normal human kidney sections (c) using FSGS-9 and FSGS-10 plasma (1:10 dilution). No overlap of IgG with PODXL was observed in paraffin sections. In frozen sections, extensive endogenous IgG background signal limited the ability to perform colocalization analysis. Scale bar: 50 μm.

### SUPPLEMENTARY FIGURE S8

## a.

**Conventional ELISA** : High background issue (this study and ref. [6] )

**Immunoprecipitation**
- IP followed by WB (this study and ref.[2,5,6]: Most reliable but less quantitative
- IP followed by ELISA (ref. [6] ): Obtain quantification and labor intensive
- IP followed by on-beads ELISA (this study, using magnetic beads): High throughput, sensitive, and quantitative; can be applied to clinical CLIA-based assays.

**Immunofluorescence**
- Staining of transfected HEK293 cells (this study)
- Staining of human kidney specimens (this study)

**IgG deposition in pathology specimens**[2,3,4]: IgG deposits appear in intracellular puncta with partial overlap with endogenous nephrin.

## b.

(1) **Tested in this study:** (Nephrin antigen types/preparations)
- a. R&D Systems His-tagged NPHS1-ECD: (Expressed from mouse cell line, **generating false-positive results**)
- b. SinoBio His-tagged NPHS1-ECD: (Expressed from HEK293 cells, **inconsistent results with FLAG-tagged**) **NPHS1-ECD for some samples**
- c. Lab-produced FLAG-tagged NPHS1-ECD: (Expressed from HEK293 stable cell line)
- d. Lab-produced full-length NPHS1: (Expressed from stable HEK293 cell line, stable cell lines lost C-terminal FLAG tag)
- e. Lab-produced FLAG-tagged full-length NPHS1: (Expressed from transient transfected HEK293)

(2) **Shirai et al.**[3] (R&D Systems His-tagged NPHS1-ECD)
(3) **Hengel et al.**[6] (Lab-produced NPHS1-ECD with His and Twin-Strep tags: Expressed from HEK293 cells)
(4) **Watts et al. and Fujita et al.** [2,7] (Lab-produced His-tagged NPHS1-ECD: Expressed from HEK293 cells)
(5) **Hattori et al.**[5] (Lab-produced full-length NPHS1 with Myc-FLAG tag: Expressed from HEK293T cells)

**Supplementary Figure S8.** (**a**) Summary of assay methods used in this study and previously published studies. (**b**) Summary of antigens used in this study and previously published studies.

### SUPPLEMENTARY TABLE S1

**Supplementary Table S1:** Summary of the study cohort.

| Case number | Diagnosis |
|---|---|
| 12 | Healthy control (Healthy Ctr) |
| 5 | Non-kidney disease transplantation control (TX Ctr) |
| 5 | IgA nephropathy (IGAN) |
| 3 | Minimal change disease (MCD) |
| 3 | Membranous nephropathy (MN) |
| 20 | Recurrent focal segmental glomerulosclerosis (rFSGS) |
| 8 | Non-recurrent focal segmental glomerulosclerosis (nrFSGS) |
| 4 | Secondary focal segmental glomerulosclerosis ($2^{nd}$ FSGS) |
| 10 | Primary focal segmental glomerulosclerosis (Primary FSGS) |
| 23 | Systemic lupus erythematosus (SLE) |
| 34 | Type I diabetes (T1D) |
